# Cycling Cadence Selections at Different Saddle Heights Minimize Muscle Activation Rather Than Energy Cost

**DOI:** 10.1101/2024.05.30.596596

**Authors:** Cristian D. Riveros-Matthey, Mark J. Connick, Glen A. Lichtwark, Timothy J. Carroll

## Abstract

Unlike walking and running, people do not consistently choose cadences that minimize energy consumption when cycling. Assuming a common objective function for all forms of locomotion, this suggests either that the neural control system relies on indirect sensorimotor cues to energetic cost that are approximately accurate during walking but not cycling, or that an alternative objective function applies that correlates with energy expenditure in walking but not cycling. This study compared how objective functions derived as proxies to 1) energy cost or 2) an avoidance of muscle fatigue predicted self-selected cycling cadences (SSC) at different saddle heights. Saddle height systematically affected SSC, with lower saddles increasing SSC and higher saddles decreasing SSC. Both fatigue-avoidance and energy-expenditure cost functions derived from muscle activation measurements showed minima that closely approximated the SSCs. By contrast, metabolic power derived from *VO*_2_ uptake was minimal at cadences well below the SSC across all saddle height variations. The mismatch between the cadence versus muscle activation and the cadence versus metabolic energy relations is likely due to additional energy costs associated with performing mechanical work at higher cadences. The results suggest that the nervous system places greater emphasis on muscle activation than on energy consumption for action selections in cycling.

## Introduction

Optimal control theory has been proposed to explain how motor behaviors are selected and produced (or “emerge”) in the face of redundancy (Bertram 2005; Ralston 1958; Hoyt 1981; Zarrugh, Todd, and Ralston 1974). This theory posits that we organize our movements to achieve specific functional goals, such as moving from one place to another, while also meeting generic performance objectives, for example using the least possible energy. Meeting these generic performance objectives can also be thought of as minimizing the costs of movement (Anderson and Pandy 2001; Bertram 2005; Donelan, Kram, and Kuo 2001; Zarrugh, Todd, and Ralston 1974; Ralston 1958; Candau et al. 1998; Cappellini et al. 2006). In walking, for example, people select stride lengths and rates that are aligned with the minimum metabolic energy consumption for any given speed (Holt, Hamill, and Andres 1990; Minetti and Alexander 1997; Umberger and Martin 2007). In general, humans and other animals tend to locomote in ways that minimize energy use, which suggests that the neural mechanisms that underlie preferences for movement economy may have an evolutionary basis (Alexander 1996). Neural control systems that minimize energy use in locomotion would have clear ethological advantages for survival, so there is considerable research effort to determine how the nervous system might implement such systems (Snaterse et al. 2011; Selinger et al. 2015; Selinger et al 2019; Wong, Selinger, and Donelan 2019).

Despite the attractiveness of energy minimization as part of a general objective function for locomotion, most people do not minimize energy consumption when selecting cadences for cycling (Brisswalter et al. 2000; Alejandro Lucia, Jesus Hoyos 2000; Marsh and Martin 1997; Brennan et al. 2019). Optimal cadences for energy economy are typically around 55-65 rpm, whereas self-selected cadences (SSC) often range between 85-95 rpm. This presents a valuable opportunity to critically assess the energy minimization phenomenon. At a minimum, it suggests that humans do not always effectively implement optimal control according to a pure energetic minimization cost function. This could indicate that an “optimal control-like” neural control system uses alternative sensorimotor signals as a proxy for energy expenditure, and that these signals do not correlate with energy use as well in cycling as they do in walking. Alternatively, it is possible that an entirely different objective function drives selection of motor activation patterns in locomotion, and that this objective function correlates well with energy use during walking and running but not in cycling. Studying cadence preferences in cycling may provide insights to address these questions.

A candidate for an energy expenditure proxy in optimal control objective functions is the degree to which muscles are activated by the central nervous system. Although an alignment between muscle activation minimization and energy expenditure minimization is well-documented in various locomotion contexts (Russell and Apatoczky 2016; Sousa and Tavares 2012), there are also examples in which muscle activation is not an accurate proxy for movement economy (Miller et al. 2012; McDonald et al. 2022; Silder, Besier, and Delp 2012; Umberger 2010; Beck et al. 2019). Importantly, muscle activation might be more straightforward for the CNS to monitor as an optimal control cost, due to the rapid availability of efference copy information (Wolpert and Flanagan 2001) and muscle afferent proprioceptive feedback (Monjo, Terrier, and Forestier 2015; Proske and Allen 2019), in contrast to slower estimates of metabolic energy expenditure that would require integration from multiple sensors of different classes (Wong, Selinger, and Donelan 2019).

Many authors have considered cost functions for locomotor control derived from muscle activation (Griffin, Roberts, and Kram 2003; Kipp, Grabowski, and Kram 2018; Kram and Taylor 1990). Such “energy proxy” or weighted activation cost functions are based on the assumption that active skeletal muscles are the primary consumers of metabolic energy during task-oriented motions, thereby dictating overall metabolic energy expenditure. In essence, such cost functions quantify the volume of activated muscle tissue used to generate force during movement. They have been shown to correlate well with energy expenditure during walking on the flat (Griffin, Roberts, and Kram 2003; Umberger 2010), but not during uphill walking (McDonald et al. 2022). Muscle volume activation was also approximately minimized close to the SSC across varying power demands in simulated cycling (C. Riveros-Matthey, Carroll, Connick, et al. 2023).

There are, however, alternatives to the hypothesis that metabolic cost (or an associated proxy) minimization drives locomotor behavior. Indeed, although metabolic cost minimization is commonly justified from a “survival strategy” perspective (Pontzer 2017), a desire to avoid muscle fatigue, which is a reduction in maximal voluntary muscle force generating capacity caused by exercise (Gandevia 2001), should not be underestimated as a potential “survival strategy” control cost. In situations such as prolonged prey pursuits or evasive action, muscle activation strategies that delay fatigue and prevent movement failure (e.g., when muscles can no longer produce sufficient force to meet task demands) become paramount well before overall energy depletion becomes relevant (Marino, Sibson, and Lieberman 2022; Pontzer 2017). Thus, objective functions that seek to avoid muscle fatigue appear plausible candidates to shape behavior in locomotion tasks.

This general perspective has led to development of “fatigue-like” or “fatigue avoidance” cost functions derived from muscle activation measurements (Ackermann and van den Bogert 2010; Miller et al. 2012; Neptune and Hull 1999). These cost functions encourage the minimization of muscle activation patterns by evenly distributing the workload among muscle groups (Ackermann and van den Bogert 2010) and penalizing higher activations to mitigate the risk of premature muscle fatigue (Ackermann and van den Bogert 2010; Miller et al. 2012). Fatigue avoidance cost functions closely align with overall energy expenditure cost functions in flat walking and consequently correlate well with behavioral selections (Russell and Apatoczky 2016; Sousa and Tavares 2012). Crucially, fatigue avoidance cost functions are also good predictors of action selection during inclined walking (McDonald et al. 2022; Silder, Besier, and Delp 2012) and experimental cycling studies (MacIntosh, Neptune, and Horton 2000; Takaishi et al. 1996; C. Riveros-Matthey, Carroll, Lichtwark, et al. 2023).

Investigating cost functions for action selections under different mechanical constraints has provided valuable insights into their significance in walking (Selinger et al. 2015; McDonald et al. 2022). Likewise, in cycling, modifications to bicycle geometry can exert influence over the energetic and mechanical landscape for action selection (Nordeen-Snyder 1977; Rankin and Neptune 2010; Peveler 2008). For example, adjusting the saddle position influences the limb range of motion (Rodrigo Rico Bini, Hume, and Kilding 2014; Rodrigo R. Bini, Tamborindeguy, and Mota 2010) and the activation and force requirements of muscles involved in work generation (Connick and Li 2013; Sanderson and Amoroso 2009; Rankin and Neptune 2010). Consequently, variations in saddle height result in distinct energetic-mechanical landscapes and, potentially, a different SSC (Rugg and Gregor 1987). Any specific features of behavior that are minimized across distinct energetic-mechanical landscapes would appear to be prime candidates for a general objective function for the neural control of locomotion.

In this study, the main goal was to test how changing saddle height affected cycling behavior. We tested whether a set of optimization criteria based on muscle activation and metabolic cost can accurately predict SSC amidst diverse mechanical cost landscapes. We used breath-by-breath gas analysis to measure metabolic power, and electromyography of the right lower limb to compute muscle activation-driven cost functions. We hypothesized that cyclists would choose different cadences in response to changes in saddle heights, and that fatigue avoidance cost functions would consistently align with SSC regardless of saddle height changes. This would be consistent with the idea that the control strategy employed by the nervous system in action selection for cycling prioritizes the avoidance of fatigue or the prevention of movement failure over the minimization of energy use.

## Material and methods

### Subjects

We estimated effect size (Cohen’s d) of 1.24 from a pilot study to detect differences in the average sum of squared normalized muscle activations between specific paired groups (saddle heights at 90 rpm) with a power of 95%, resulting in the required eleven participants as a sample size. We recruited twelve individuals to account for the potential loss of data. Specifically, we enrolled male cyclists who met the criteria of being daily riders at levels 3-4, indicating their considerable experience and involvement in the discipline from the university community and cycling sports clubs. Before completing the experimental trials, the participants’ age, mass, and height (mean and standard deviation, age= 34.8 ± 9.5 yr., mass= 79.1 ± 10.2 kg, height= 1.78 ± 0.08 m) were measured using a measuring tape and scales, respectively. This study was approved by the Human Research Ethics Committees (HRECs) of the University of Queensland (2022/HE001934) and participants provided written informed consent.

Cycling was performed entirely on a bicycle ergometer (Excalibur Sport, Lode BV, Groningen, The Netherlands). This ergometer was adjusted to each subject’s anthropometric characteristics, with special attention paid to the handlebars and saddle. The saddle height was normalized to 100% greater trochanter height (h_t_) (Rodrigo Rico Bini et al. 2014). The trunk angle was standardized between 35 and 45°, depending on each subject’s preference, with their hands positioned on the handlebar drops. The trunk angle was defined through the horizontal line with respect to the line that connects the anatomical landmarks of the acromion and greater trochanter. Visual feedback was used to ensure that posture was maintained during the experimental sessions. The crank length was set to 175 mm and participants wore clothes cleated shoes clipped in the pedals (SH-R070, Shimano, Osaka, Japan; SH-R540, Shimano, Osaka, Japan).

### Experimental design

Participants took part in three days of assessments, with each experimental session lasting less than 2 hours. The first day session comprised a series of maximal voluntary contraction (MVC) tests that aimed to obtain the maximum EMG activation in isometric conditions for all muscles sampled. The participant performed a 5s maximal voluntary contraction of each muscle group against a resistance provided by the researcher. Participants were encouraged to do their best at each trial. Then, after a 5min warm-up of 90rpm at 100W, participants completed two SSC assessment conditions separated by an exploration protocol. Each SSC assessment was performed over 3minutes. The exploration condition required participants to cycle for 30s at cadences of 50, 65, 80, 95, 110 rpm at a fixed power output (2.5 *W · kg*^-1^), in a randomized order. The SSC was defined as the mode of the instantaneous cadences over the last two minutes for each saddle height condition. Afterwards, a protocol to measure the metabolic, muscle activation and mechanical characteristics of cycling at different cadences took place. Riders cycled at the same power output with five different cadences (50, 65, 80, 95 and 110rpm) for 3 minutes each. There was 3 minutes rest between trials (see Figure 1). The following measurements were taken during each trial: marker-based 3D motion capture, surface electromyography, breath-by-breath gas analysis, radial tangential pedal force, and crank angle on the cycle ergometer.

**Figure 1:**
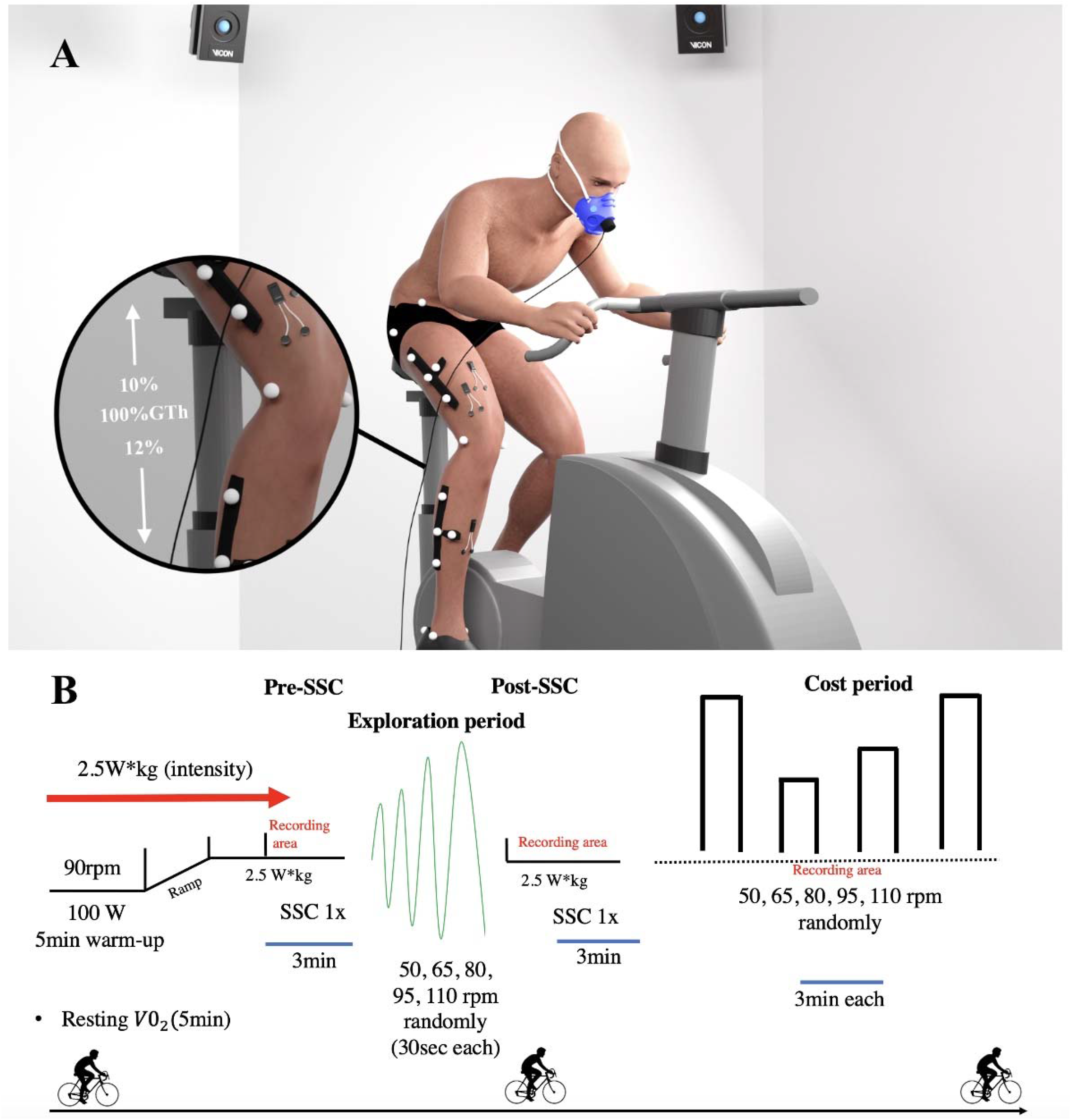
Schematic representation of the experimental design performed for each saddle height variation (normal saddle: 100% of the greater trochanter ( ); low saddle and high saddle: 12% lower and 10% higher than the normal variation, respectively). Data were extracted from 3-dimensional motion capture, surface electromyography, breath-by-breath gas analysis and radial tangential force on the cycle ergometer (A). The experimental protocol was applied to each subject in the three days of assessment, each testing a different saddle height variation (B).

The second and third-day assessments involved a similar protocol to the first-day assessment. However, a targeted mechanical perturbation was added. On the second day, we implemented a low saddle condition, adjusting the saddle height to 88% of the height measured from h_t_during the first day session. On the third day, we introduced the high saddle condition, setting the saddle height to 110% of Th. The mean trunk angle remained consistent across saddle conditions.

### Instruments

#### Electromyography (EMG)

EMG electrodes were placed on the gluteus maximus (GMAX), vastus lateralis (VL), rectus femoris (RF), biceps femoris (BF), semitendinosus (ST), tibialis anterior (TA), soleus (SOL) and gastrocnemius medialis (GM) muscles of the right leg. Wireless amplifiers were connected to electrodes and used to collect surface EMG data (Cometa, Milan, Italy) at 2kHz, synchronously recorded with motion capture data, using the A/D board under the control of the Nexus 2.12 Software. The electrodes were placed on the subject’s skin with a 2 cm interelectrode distance according to SENIAM guidelines (Hermens et al. 2000). Prior to application of the electrodes, hair over the muscle was shaved with a disposable razor before the placement and the skin was cleaned with 70% isopropyl alcohol.

#### Metabolic Cost

Metabolic data were collected using a portable breath-by-breath gas analyzer (Metamax 3B, CORTEX BiophysikGmbH, Germany). During each trial, the exchange of *V0*_2_ and *VC0*_2_ was continuously measured. Before each test, the gas analyser was calibrated considering the atmospheric pressure and environmental oxygen concentrations. A 5-minute resting *V0*_2_ was considered before each collection, while the steady-state was calculated at the last minute of each cycle ergometer trial when the *V0*_2_ variation was relatively constant (less than 10% of variation).

#### Kinematics

Participants’ joint movements were recorded using six video-based 3-dimensional motion-capture cameras (Vicon, UK) and software (Nexus 2.12, Vicon, UK). On the participants’ right leg skin, twenty-two small light-weight reflective markers were attached. Rigid four-marker clusters were placed on the mid-thigh and mid-shank, markers were also placed on the anterior and posterior iliac spines, greater trochanter, medial and lateral epicondyles, medial and lateral malleoli, calcaneus, and the first and fifth metatarsal heads individually. A trial was captured in a static standing position with their arms crossed, after placing the reflective markers on each cyclist. The static recording was then applied to data analysis for scaling purposes. Reflective markers were positioned on the back of the saddle and at the left and right rear support angles of the cycle ergometer to establish a bicycle coordinate system to which all marker coordinates and force could be referenced. The cameras tracked the 3D trajectories from each marker at 100Hz.

#### External forces

Instrumented cranks (Axis, SWIFT Performance, Brisbane, Australia) measured the orthogonal crank forces, crank torque (tangential force * length) and the crank axial force (radial force) when a participant was pedaling. Since the crank angle was also recorded simultaneously, the vertical and horizontal components of pedal forces Fz and Fx were derived. These cranks were represented in the bicycle coordinate system. All data were sent via digital signal to a radio receiver, converted to from digital to analog and recorded at 100 Hz, synchronously with all motion capture and EMG data via an A/D board controlled by the Nexus 2.12 Software. A static calibration was performed on the instrumented cranks before each collection.

### Data Analysis

#### EMG analysis and cost functions

EMG signals were processed using a 15/500 Hz bandpass filter to eliminate non-physiological noise from the signal. To account for any analog channel offset, the median EMG signal of each muscle was subtracted from the signal. The RMS was calculated with a moving window width of 50ms as an index of the signal amplitude. For each muscle and task, we then quantified mean cycle activation 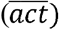 by averaging EMG-RMS signals over the task cycle 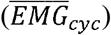 and dividing by the maximum RMS envelope values during the maximal voluntary contraction (*EMG*_*mvc*_):

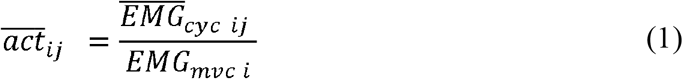

where the subscript *j* corresponds to the cycle number relative to 10 cycles recorded (*j* = 1,2,3, … 10), and *i* corresponds to the muscle of interest (*i* = 1, 2, 3, … 8).

We also quantified peak cycle activation (*act*_*peak*_) by taking the maximum of the enveloped EMG-RMS signal across the cycle *(EMG*_*peak*_) and dividing by *EMG*_*mve*_ :

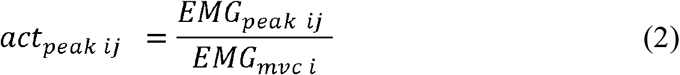

We derived a family of cost functions based on EMG, modified from the generic cost function (*CF*) proposed by Ackermann & van den Bogert (2010).

#### Fatigue avoidance cost function

The sum of squared normalized muscle activation 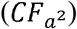 has classically represented a fatigue avoidance cost function (Miller et al. 2012; Ackermann and van den Bogert 2010). This approach, characterized by employing exponents (*p*) greater than 1, penalizes significant muscle activations that lead to fatigue, irrespective of muscle size (Ackermann and van den Bogert 2010). Nevertheless, for a more robust fatigue avoidance effect, we also considered a refined version of this cost function that incorporates the square of the peak activation observed within each cycle (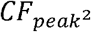 ; Eq. 3), rather than the average. We posit that minimizing peak activation squared should prioritize the prevention of muscles from operating at near maximum activation levels.

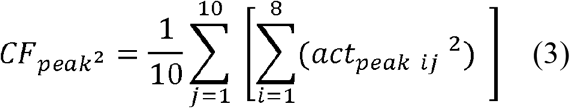

#### Minmax fatigue avoidance cost function

The traditional minmax cost function is a variation of the fatigue avoidance class of cost functions that focuses on the activity of the most strongly activated muscle. The cost function treats the maximally activated muscle as the control parameter and ignores the activity of other muscles (Rasmussen, Damsgaard, and Voigt 2001). This criterion operates under the assumption that task performance is restricted by the “weakest link” among the involved muscles, so muscle coordination should prioritize the prevention of force production failure in any individual muscle rather than reducing activation on average across muscles. We modified the traditional function to emphasize and penalize higher activations by including the square of the peak activation of the most strongly activated muscle (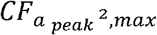; Eq. 4).

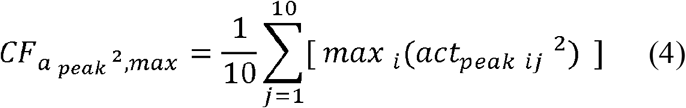

#### Weighted activation “energy proxy” cost function

The sum of the volume-weighted muscle activations was considered as a viable cost function to reflect a muscle activity-based proxy of energetic cost. This cost function weights the contributions of each muscle’s activation level according to its size (Miller et al. 2012; Kram and Taylor 1990; Biewener et al. 2004), and will therefore reflect the total amount of active muscle better than cost functions that are not volume-weighted. The normalized activation (*act*_*mode*_) for each muscle was weighted (*ω*_*i*_= *vol*) (McDonald et al. 2022) based on the muscle volumes extracted from Handfield et al. (Handsfield et al. 2014). Given that this cost function *CF*_*a*,*vol*_ (Eq. 5) reflects both muscle activation and size, we adhered to the conventional approach, where the normalized activation of muscles is averaged (*act*_*µ*_) and weighted by volume (*ω*_*i*_= *vol*) with a singular exponent (*p* =1).

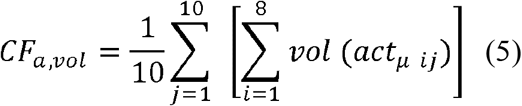

#### Metabolic power cost function

A metabolic power cost function (*CF*_*mp*_) was computed from expired oxygen (*V0*_2_) and carbon dioxide (*C0*_2_) measurements from the last minute of each cycle ergometer trial. The *CF*_*mp*_ for each condition was calculated according to standard equations (Kipp, Byrnes, and Kram 2018) based on *V0*_2_ and *VC0*_2_ uptake.

#### Joint-specific powers and VL MTU analysis

The method for determining joint-specific powers is detailed in Riveros-Matthey et al. (C. Riveros-Matthey, Carroll, Lichtwark, et al. 2023). Briefly, data obtained from the motion capture markers, force and angle of the instrumented pedals were processed in MATLAB (R2019a, Mathworks Inc., USA). Force components were transformed from the crank position to the bicycle coordinate system. Subsequently, the kinematic and kinetic data were represented in a rigid body system using the OpenSim software (inverse kinematics and inverse dynamics tools) to obtain joint-specific powers, which are the product of angular velocity and joint moments (Elmer et al. 2011). The OpenSim muscle analysis tool was used to determine the lengths and velocities of the VL MTU under each condition.

### Statistical analysis

A repeated measures one-way ANOVA was used to test the main effect of the saddle heights on SSC. Muscle activation-driven cost functions and metabolic power were fitted using second-order polynomial regressions to assess their relationship with cadences in each saddle height condition. The correspondence between the cadence that minimized each cost function (predicted by the nonlinear regression) and the experimentally measured SSC across all saddle height conditions was then assessed using linear mixed effects models (LMEM). In these models, the experimental SSC was treated as the response variable and the cadence at which each cost function was minimized was treated as a fixed effect, while subjects were included as a random effect to account for individual variations in *slope* and y-intercept. In the LMEM approach, a *R*^2^ and *slope* were calculated to assess the correspondence between the experimental and predicted metrics. Correlations were interpreted based on the magnitude of *R*^2^ values: ≤ 0.29 as weak, 0.30 to 0.39 as moderate, 0.40 to 0.69 as strong, and ≥ 0.70 as very strong. Additionally, we used a two-way repeated measures ANOVA to assess the absolute agreement between the cost function minimum cadences and the SSC, a benchmark to establish which of the costs may drive action selections in cycling. These scores were derived by computing the absolute differences between the SSC and the cadence at which each cost function reached its minimum across subjects and various saddle heights. We considered the main effects of saddle heights and cost function form and their interactions on cost function difference scores. For all tests, an alfa level of 0.05 was used. Prism 7 (GraphPad Software Inc., La Jolla, CA) and MATLAB (R2019a, Mathworks Inc., USA) were used to run all statistical tests.

## Results

Participants cycled at a power output of 2.5 *W* · *kg*^-1^ (197 ± 25W) with three saddle heights (normal saddle: 100% of Th, 94.7 ± 4 cm; low saddle: 88% of Th, 83.3 ± 3 cm; high saddle: 110% of Th, 104 ± 5 cm). Average *V0*_2_ consumption rates were 2.98 ± 0.2, 3.10 ± 0.3, and 3.29 ± 0.3 L.min^-1^ for normal, low, and high saddle conditions, respectively.

### SSC responses across saddle heights

Figure 2 displays the cadences that were self-selected as preferable before and after the exploration period. There was a main effect of saddle position on pre-(*P=*0.001) and post-exploration (*P <* 0.001) SSCs, with a general trend toward lower cadences with higher seat heights (pre - low: 94.9 ± 10 rpm, normal: 89.7 ± 9 rpm; high: 81.6 ± 11 rpm; Figure 2A; post - low: 101.9 ± 8 rpm; high: 78.3 ± 13 rpm; Figure 2B). At both measurement occasions, SSC was significantly lower for high than low saddle conditions (both: *P =* 0.001). However, after exploration, SSC was only significantly lower for the high saddle position compared to the normal saddle (*P =* 0.001; Figure 2B). There was no significant difference between normal and low saddles (before: *P =* 0.30; after: *P =* 0.24). We posit that post-exploration results should more accurately depict a low-level neural action selection process than pre-exploration findings given the increased exposure to perturbations and the greater consistency in the effect of saddle heights observed across participants. Consequently, subsequent analyses exclusively focused on the post-exploration period for muscle activation-driven cost functions and metabolic power.

**Figure 2:**
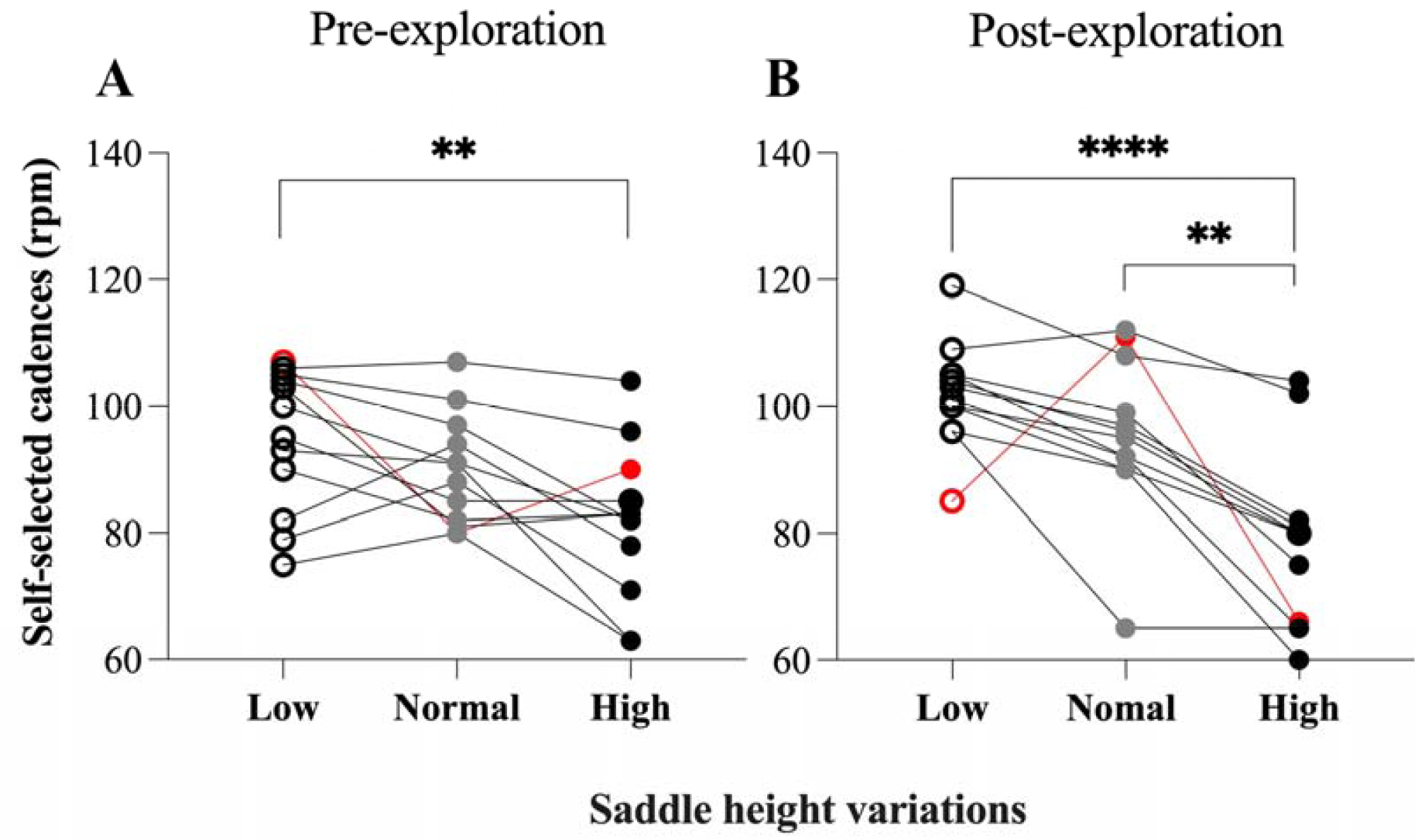
SSC pre (A) and post (B) exploration period at normal, low, and high saddles across subjects. The highlighted figures and lines represent the average of the subjects. Red dots and lines correspond to the excluded participant.

Moreover, our study primarily aimed to understand the relationship between the cycling cadences that people naturally choose and various muscle activation and energetic costs. The conclusions rely upon the assumption that participant choices reflect neural processes for action selection that determine motor behavior based on low-level sensorimotor variables. However, human motor behavior is also subject to high-level strategic decisions that may not reflect the low-level neural processes that typically guide subconscious action selection. Such strategic decisions may lead to idiosyncratic responses to experimental manipulations and would obscure relationships between sensorimotor cost variables and action selection processes. One participant in this study showed choice behavior indicative of such idiosyncratic high-level decision-making. Their cadence selections varied dramatically between the pre- and post-exploration measurements of SSC and showed a pattern across seat height variations that was fundamentally distinct from all other members of the sample (see red data points in figure 2). Thus, we excluded the data from this participant from regression and ANOVA fits comparing SSC to cost function predictions. The final analysis therefore involved eleven datasets for muscle activation-driven cost functions and metabolic power.

### Muscle activation-driven cost functions

#### Fatigue avoidance cost function

The relationship between cadence and peak activation squared 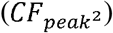 displayed a curvilinear relationship in each saddle position, with greater 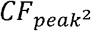 values at high and low cadences in the normal and low saddle conditions. Conversely, at the highest saddle, pronounced costs were evident at higher cadences, specifically at 110rpm (Figure 3A). The comparison between predicted minima and the SSC via linear mixed effects model revealed a very strong correlation (*R*^2^=0.77) for 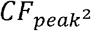 . Furthermore, the models revealed that the regression *slope* was 1.21 (Figure 3B).

**Figure 3:**
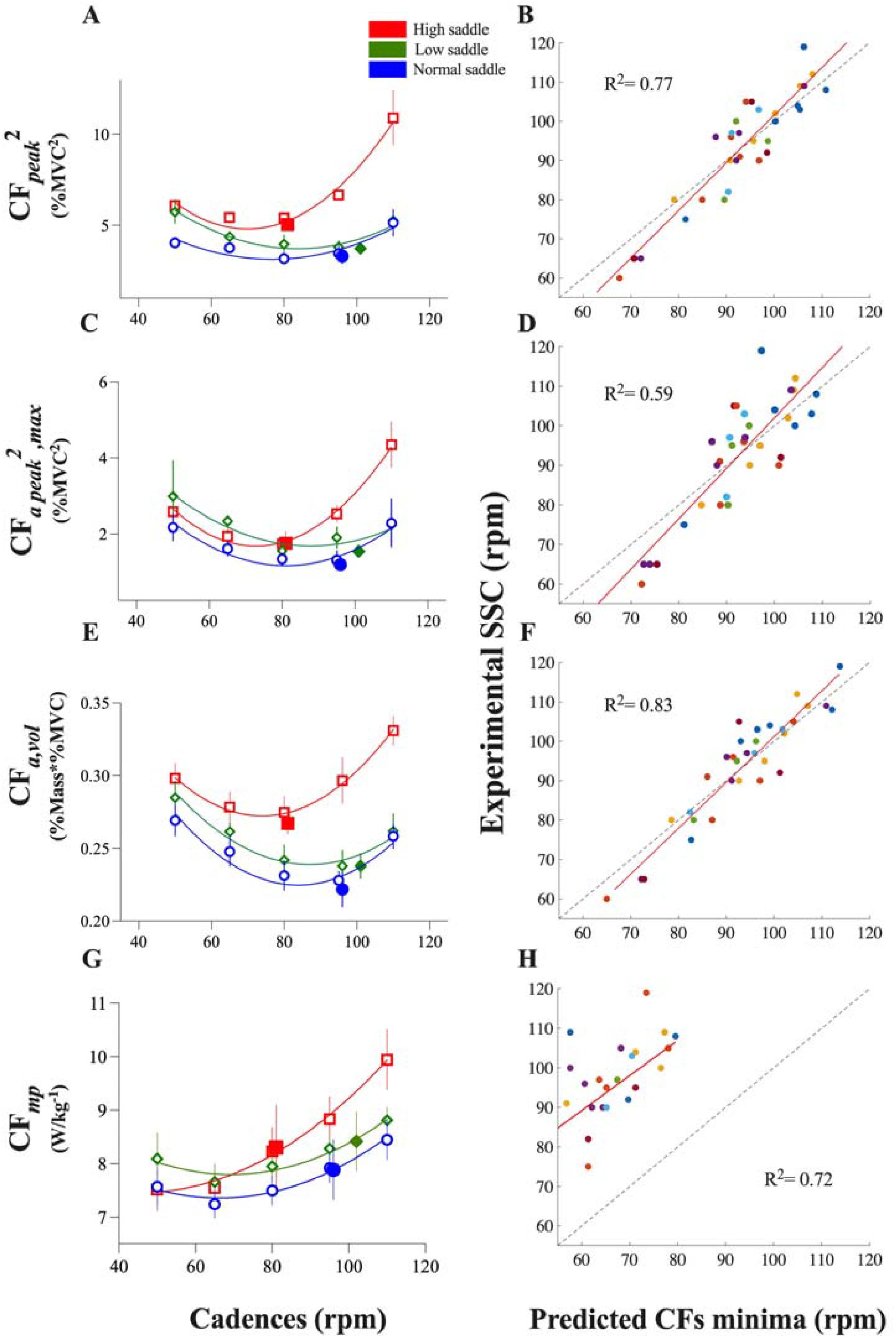
A second-order polynomial non-linear regression (left) and linear mixed effects model (LMEM) (right) (from A to H) analysis on: 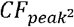 (A and B), 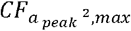 (C-D), *CF*_*a*,*vol*_ (E-F) and *CF*_*mp*_ (G-H) cost functions at 50, 65, 80, 95, 110 rpm and SSC post-exploration across saddle heights. The SSC at the post-exploration is 95.6 ± 10 rpm at the normal saddle (filled circle), 102.3 ± 9 rpm at the low saddle (filled diamond) and 79.4 ± 11 rpm at high saddle variation (filled square). Data from the non-linear regression charts are means ± SD. To address variability in muscle activation-driven cost functions, a 95% confidence interval bootstrap technique was used to enhance the visualization of the relationship between costs and cadences from nonlinear regressions. Confidence limits were based on 100 resampled data points. Nonlinear regression was computed for each set of cost function data, including cadences and saddle heights across subjects, while the full dataset was used for statistical analysis. Blue, green, and red figures and lines correspond to normal, low, and high saddle heights, respectively. Subjects are represented by colored dots on the LMEM charts.

#### Minmax fatigue avoidance cost function

A curvilinear relationship between cadence and the *minmax* derived cost function was observed across saddles, with consistent increases in cost at low cadences (50rpm) in all saddle positions (Figure 3C). In contrast, the highest costs were observed at higher cadences during the high saddle position (Figure 3C). The linear mixed effects model showed a strong correspondence (*R*^2^=0.59) between the SSC and 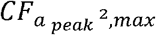 cadence minima, with a *slope* of 1.27 (Figure 3D).

#### Weighted activation “energy proxy” cost function

The relationship between *CF*_*a*,*vol*_ and cadence displayed a curvilinear pattern with a ‘U-shaped’ trend across all saddle positions. Notably, the high saddle variation displayed higher costs across cadences (Figure 3E). The linear mixed effects model showed a very strong correlation (*R*^2^=0.83; Figure 3F) between the cadence in which the predicted volume-weighted cost functions minima were observed and the experimental SSC. Furthermore, the cost function regression model line reached a *slope* of 1.16.

#### Metabolic power cost function

There was a non-linear relationship between the *CF*_*mp*_ cost function and cadence across all saddle heights, with a minimum that varied according to the saddle height performed. For normal saddle conditions, the minimum was reached at 66 ± 7 rpm (Figure 3G). The low and high saddles showed an inverse relationship where the low saddle resulted in a minimum metabolic power at a higher cadence of 69 ± 8rpm, while the high saddle variation resulted in a minimum metabolic power at cadences near 50 ± 4rpm (Figure 3G). Although a very strong ( : 0.72; Figure 3H) correspondence was found between the cadence of minimum metabolic power and the SSC ( = 0.91), the intercept of the linear fit was considerably offset from zero, such that the cadence of minimum energetic cost consistently underestimated the SSC.

### Degree of closeness between the cost functions and the SSC

We calculated the absolute difference between the cadences that minimized each cost function and the SSC for each participant at each seat height to compare how closely the minimum cadence in each cost function matched to SSC. The two-way ANOVA revealed a main effect of saddle height conditions (*F*_(1,12)_= 4.5, *P*=0.045, η^2^ = 0.05) and cost functions (*F*_(2,23)_= 48, *P*< 0.001, η^2^ = 0.42) on difference scores, but no interaction effect (saddle height conditions x cost functions) (*F*_(2,29)_= 2.5, *P*=0.076, η^2^ = 0.04). The muscle activation-driven cost functions were not significantly different from each other (e.g. 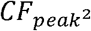 : degree of closeness of ∼15rpm and *CF*_*a*,*vol*_ a degree of closeness of ∼9.5rpm). Moreover, the *CF*_*mp*_ difference (∼33.2rpm; Figure 4) from SSC was significantly greater than all three muscle activation-driven cost functions (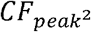 : *P*<0.001, degree of closeness =∼15rpm; *CF*_*a*,*vol*_:*P*<0.001, degree of closeness = ∼9.5rpm, and 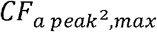 : *P*<0.001, degree of closeness = ∼16.3 rpm).

**Figure 4:**
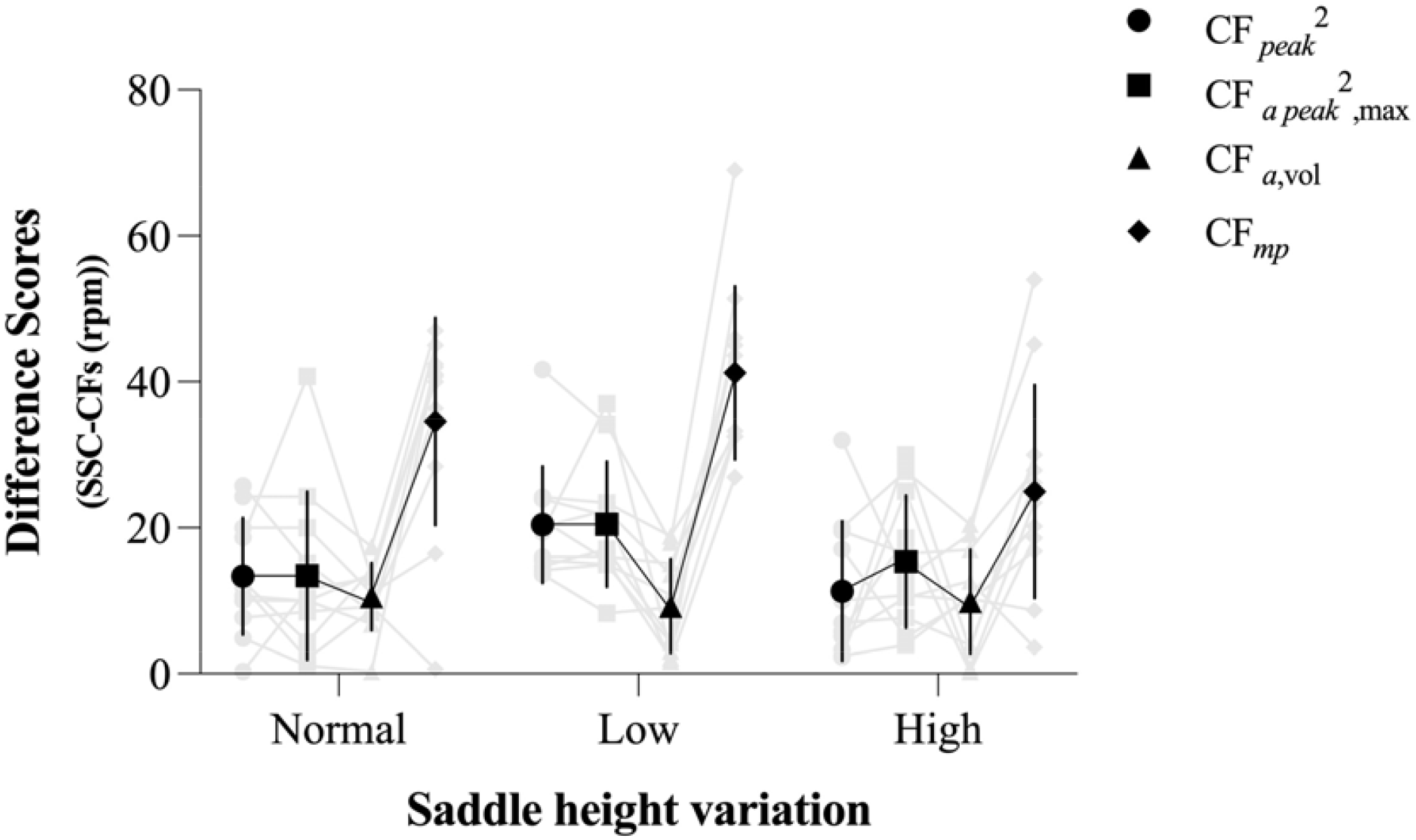
Difference scores among cost functions over saddle heights. Mean ± SD , , and cost functions across normal, low, and high saddles. Two-way ANOVA, considering an αlevel of 0.05.

## Discussion

The goal of this study was to test how well cost functions based on muscle activation and metabolic cost can predict the cycling cadences that people select across saddle heights. In line with our first hypothesis, changes in saddle height led to changes in SSC. Specifically, the SSC tended to increase as saddle height was reduced. Our study also showed that the minima of all three-muscle activation-driven cost functions fell at cadences close to the SSCs across all saddle heights. In line with previous findings (Coast and Welch 1985; Marsh and Martin 1993), the minimum metabolic power occurred at cadences much lower than the SSC, confirming that humans do not choose to minimize whole-body energy expenditure during cycling, irrespective of the mechanical consequences of seat height manipulations.

A key finding of our study is that the saddle heights elicited changes in the cadence selected by cyclists, both before and after the exploration period. After the exploration period, cyclists selected higher cadences of approximately ∼103rpm (Figure 2B) while the saddle height was low. Conversely, at a higher saddle height, the cadence selected dropped to values around ∼80.5rpm (Figure 2B). The variations in the SSC might be linked to changes in kinematics, such as knee angles, thereby impacting the force-length relationship of the extensor muscle fibers, particularly in the VL muscle (Rankin and Neptune 2010). These alterations were indirectly observed through the analysis of knee structures, highlighting a shift in operating length relative to the optimum (supplementary Figure S1D), muscle activation (supplementary Figure S1A), and force requirements (supplementary Figure S1B-C) across different saddle heights. Specifically, the VL muscle operates at longer, less optimal lengths at lower saddle heights and vice versa at higher saddle heights (Rankin and Neptune 2010; supplementary Figure S1D). A preference for higher cadences at low saddles is likely due to a reduction in the necessary knee extensor moment (supplementary Figure S1C) compared to high resistance / low cadence conditions (Rugg and Gregor 1987). By contrast, in high saddle conditions, cyclists prefer lower cadences due to the significant penalty associated with maintaining precise control over force projection on the pedal at higher cadences (Rodrigo R. Bini, Tamborindeguy, and Mota 2010; Kruschewsky et al. 2018). The increased distance between the saddle and the pedal causes this penalty, which reduces the cyclist’s mechanical advantage when pushing on the pedal (Leavitt and Vincent 2016). Thus, these kinematic changes and adjustments in the operating lengths of the muscle fibers represented new metabolic-mechanical landscapes, providing an opportune scenario to reveal what criteria drive behavior during cycling.

Unsurprisingly, cyclists expended more energy at higher and lower saddle heights than normal. Raising the saddle height had the greatest effect on metabolic power (>8% increase) (Figure 3G). The increase in energy rate is likely related to the additional costs of activating muscle for force production because there were similar increases on *CF*_*a*,*vol*_ (Figures 3E). However, the cadence for minimizing metabolic power was consistently lower (∼66, 69, and 50rpm for the normal, low, and high saddle, respectively) than that which minimized peak activation or the volume of active muscle (e.g., *CF*_*a*,*vol*_: ∼88, 96, and 75rpm for the normal, low, and high saddle; Figure 3E). This mismatch is most likely due to the higher energy rate required to generate work at higher muscle shortening rates (Barclay and Curtin 2023; Fenn 1923; Umberger 2010), such as those corresponding to the cadences of muscle activation-driven cost functions. Thus, if the activation cost functions influence the SSC, this may imply that the nervous system places less emphasis on optimizing energy-related factors, particularly the additional costs associated with performing positive mechanical work. Whilst weighted muscle activation might be considered a proxy for energy consumption under some conditions (e.g. walking), during cycling tasks it was clear that there was mismatch between activation and energy consumption as cadence changes, regardless of changes in energy consumption with saddle height changes.

Power-generation tasks such as cycling, in which positive work dominates, provide an ideal scenario to dissociate muscle activation-driven cost functions from measured energy expenditure. Based on our findings, we can speculate that the nervous system appears insensitive to energy expenditure incurred by doing work/actively shortening (Fenn 1924) in its control scheme. Similar activation-driven gait selection has been observed during when people were asked to choose between walking with a crouched posture or on an incline, where a dissociation between muscle activation and metabolic power minima also occurs (McDonald et al. 2022). This is consistent with the idea that activation-based control signaling might be a fundamental factor in regulating locomotor cadence in general. Whether the preference to minimize muscle activation reflects a desire to avoid fatigue or to minimize an (inaccurate) proxy for energy consumption is less clear, however. As was the case in the walking study of McDonald et al (2022), we found here in cycling that comparisons between volume-weighted muscle and non-volume weighted activation cost functions were inconclusive. Regardless of saddle height, all muscle activation-driven cost functions displayed a curvilinear relationship with cadence, and there was broad correspondence between the minima of these relationships and the SSC (Figure 3). Thus, both energy-proxy minimization and fatigue avoidance remain plausible objectives to drive the nervous system’s control scheme during locomotion.

### Limitations

Our results should be interpreted considering certain limitations. We acknowledge the inherent limitations of EMG use, such as crosstalk, skin impedance, and cancellation artifacts (Farina, Merletti, and Enoka 2014). Further, using only eight lower extremity muscles may not provide a complete picture of neuromuscular control, which could miss the participation of other important muscles. Additionally, our protocol included three days of EMG measurement, and day-to-day variability in muscle activation patterns can affect data consistency. To partially counteract this, in addition to strictly following the placement protocol, we applied marks on the skin, took photographs and made detailed notes to guarantee uniform electrode placement on each session day. Finally, the use of a limited set of eight muscles may also have reduced resolution in differentiating between the cost functions based on EMG. This limitation could have obscured fine distinctions between the functions, potentially leading to the perceived similarity observed among them.

## Conclusion

This study compared the ability of EMG-based and metabolic cost objective functions to predict cadence selections during submaximal cycling with different saddle heights. Both “energy-proxy minimization” and “fatigue avoidance” cost functions derived from muscle activation measurements effectively predicted SSC across saddle heights. Although the metabolic power curve shifted similarly to muscle activation-driven cost function curves across saddle heights, the metabolic curve minima were consistently much lower than the SSC. This suggests that whole-body energy expenditure is not the primary determinant in adjusting cycling behavior through saddle height changes. Instead, cyclists consistently minimized muscle activation-derived costs, highlighting the importance of muscle activation-based proxies for energy expenditure or fatigue avoidance when selecting a cadence.

## Declarations

## Acknowledgements

We acknowledge The University of Queensland for support and provision of resources that were essential for conducting this study.

## Ethical Statement

All participants provided written informed consent. The Human Research Ethics Committees (HRECs) of the University of Queensland (2022/HE001934) approved this study in accordance with the Helsinki Declaration.

## Data Accessibility

Research data supporting this publication are available from the UQ eSpace (C. D. Riveros-Matthey et al. 2024) repository located at https://doi.org/10.48610/6413ffe

## Authors’ contributions

C.R-M.: Conceptualization, acquisition, formal analysis, investigation, visualization, and writing of the original draft. T.J.C. conceptualized and interpreted the data, revised the manuscript, and approved the final version to be published. M.J.C. conceptualized, analyzed, and reviewed the study. G.A.L. conceptualized, supervised, and revised the manuscript, and approved the final version to be published.

All authors agreed to be held accountable for the work performed and gave final approval for publication.

## Supplementary figure

**Figure S1:**
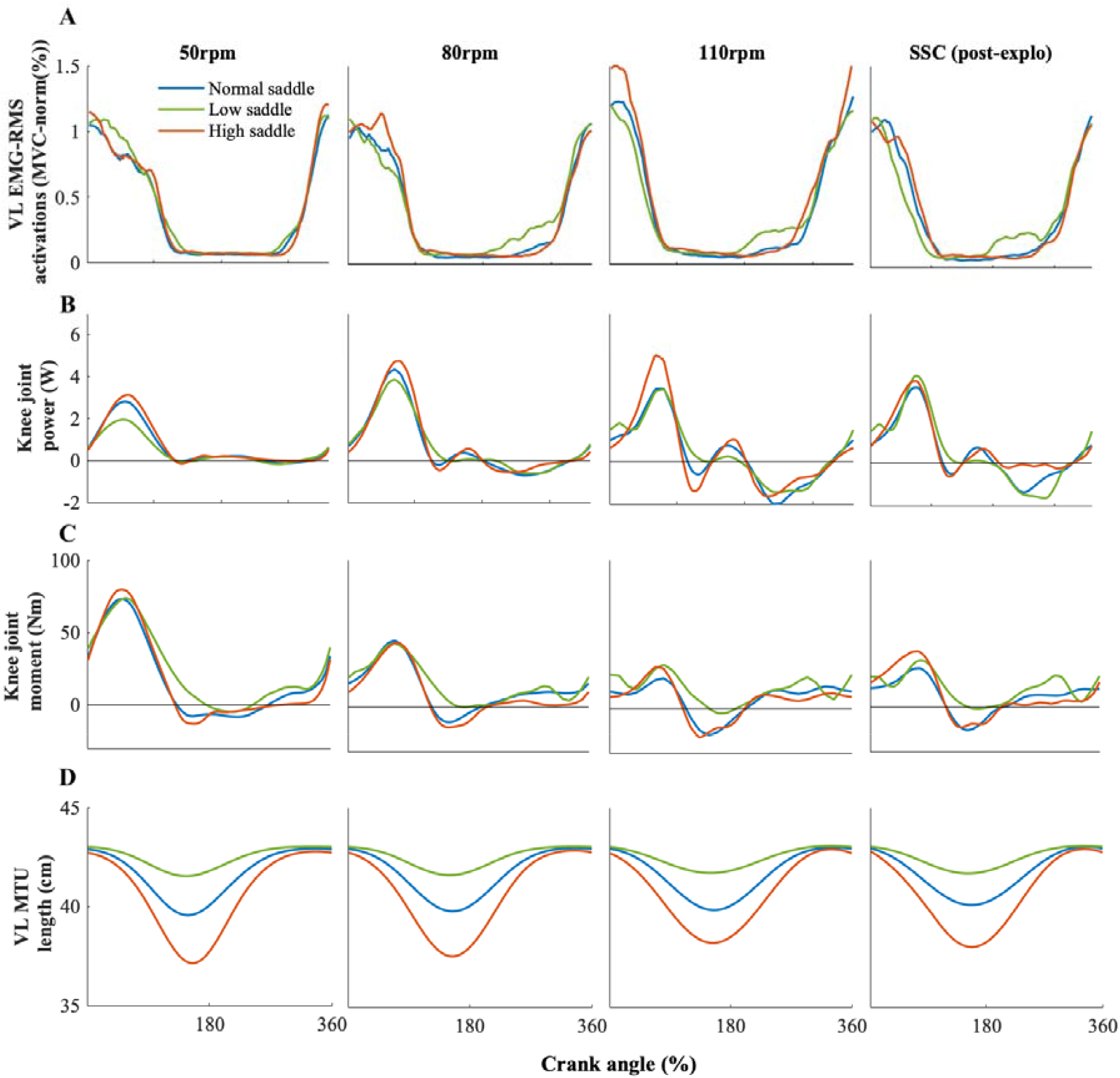
Mean waveforms of VL EMG-RMS activation (A), knee joint power (B) and moment (C) and VL MTU length (D). All the waveforms were plotted at 50, 80, 110 and SSC post-exploration period across saddle height variations over the crank angle cycle. The crank angle is 0 at top-dead-center. The SD and remaining cadences (65, 95, and SSC pre-exploration) were omitted for clarity. ANOVA approach was also used to test the main effects of saddle height variations and cadences on knee joint powers and moments. There was a main effect of saddle height conditions (knee joint power: = 38, *P* < 0.001, 0.27; knee joint moment: = 8, *P* = 0.003, 0.04), cadences (knee joint power: = 11, *P* < 0.001, 0.12; knee joint moment: = 64, *P* < 0.001, 0.54), and their interactions (knee joint power: = 2.9, *P* = 0.049, 0.05; knee joint moment: = 2.8, *P* = 0.048, 0.04) on knee positive extension joint power and moment. While knee positive extension joint power increased when the saddle variation shifted from normal to low (∼*P* = 0.01) and high (∼*P* = 0.009) conditions across cadences, a significant increase in power only was observed at 110rpm (*P* = 0.03) when transitioning from low to high saddle variation. Similarly, the knee positive extension joint moment exhibited an increase when the saddle was lowered (∼*P* = 0.02). However, during the transition from a low to a high saddle condition, an increase in knee moment was only observed at 80 rpm (*P* = 0.02) and 110 rpm (*P* < 0.001).

## References

Ackermann, Marko, and Antonie J. van den Bogert. 2010. “Optimality Principles for Model-Based Prediction of Human Gait.” Journal of Biomechanics 43 (43): 1055–60. 10.1016/j.jbiomech.2009.12.012.

Alejandro Lucia, Jesus Hoyos, and Jose L. Chicharro. 2000. “Preferred Pedalling Cadence in Professional Cycling.” MEDICINE & SCIENCE IN SPORTS & EXERCISE 13 (13): 1361–66.

Alexander, R McNeill. 1996. Optima for Animals / R. McNeil Alexander. Optima for Animals. Rev. ed. Princeton, NJ: Princeton University Press.

Anderson, F. C., and M. G. Pandy. 2001. “Dynamic Optimization of Human Walking.” Journal of Biomechanical Engineering 123 (123): 381–90. 10.1115/1.1392310.

Barclay, C. J., and N. A. Curtin. 2023. “Advances in Understanding the Energetics of Muscle Contraction.” Journal of Biomechanics 156 (May). 10.1016/j.jbiomech.2023.111669.

Beck, Owen N., Laksh Kumar Punith, Richard W. Nuckols, and Gregory S. Sawicki. 2019. “Exoskeletons Improve Locomotion Economy by Reducing Active Muscle Volume.” Exercise and Sport Sciences Reviews 47 (47): 237–45. 10.1249/JES.0000000000000204.

Bertram, J. E A. 2005. “Constrained Optimization in Human Walking: Cost Minimization and Gait Plasticity.” Journal of Experimental Biology 208 (208): 979–91. 10.1242/jeb.01498.

Biewener, Andrew A., Claire T. Farley, Thomas J. Roberts, and Marco Temaner. 2004. “Muscle Mechanical Advantage of Human Walking and Running: Implications for Energy Cost.” Journal of Applied Physiology 97 (97): 2266–74. 10.1152/japplphysiol.00003.2004.

Bini, Rodrigo R., Aline C. Tamborindeguy, and Carlos B. Mota. 2010. “Effects of Saddle Height, Pedaling Cadence, and Workload on Joint Kinetics and Kinematics during Cycling.” Journal of Sport Rehabilitation 19 (19): 301–14. 10.1123/jsr.19.3.301.

Bini, Rodrigo Rico, Patria A. Hume, and Andrew E. Kilding. 2014. “Saddle Height Effects on Pedal Forces, Joint Mechanical Work and Kinematics of Cyclists and Triathletes.” European Journal of Sport Science 14 (14): 44–52. 10.1080/17461391.2012.725105.

Bini, Rodrigo Rico, Patria A. Hume, Fabio J. Lanferdini, and Marco A. Vaz. 2014. “Effects of Body Positions on the Saddle on Pedalling Technique for Cyclists and Triathletes.” European Journal of Sport Science 14 (SUPPL.1): 37–41. 10.1080/17461391.2012.708792.

Brennan, Scott F., Andrew G. Cresswell, Dominic J. Farris, and Glen A. Lichtwark. 2019. “The Effect of Cadence on the Mechanics and Energetics of Constant Power Cycling.” Medicine and Science in Sports and Exercise 51 (51): 941–50. 10.1249/MSS.0000000000001863.

Brisswalter, J., C. Hausswirth, D. Smith, F. Vercruyssen, and J. M. Vallier. 2000. “Energetically Optimal Cadence vs. Freely-Chosen Cadence during Cycling: Effect of Exercise Duration.” International Journal of Sports Medicine 21 (21): 60–64. 10.1055/s-2000-8857.

Candau, R., A. Belli, G. Y. Millet, D. Georges, B. Barbier, and J. D. Rouillon. 1998. “Energy Cost and Running Mechanics during a Treadmill Run to Voluntary Exhaustion in Humans.” European Journal of Applied Physiology and Occupational Physiology 77 (77): 479–85. 10.1007/s004210050363.

Cappellini, G., Y. P. Ivanenko, R. E. Poppele, and F. Lacquaniti. 2006. “Motor Patterns in Human Walking and Running.” Journal of Neurophysiology 95 (95): 3426–37. 10.1152/jn.00081.2006.

Coast, J. Richard, and Hugh G. Welch. 1985. “Linear Increase in Optimal Pedal Rate with Increased Power Output in Cycle Ergometry.” European Journal of Applied Physiology and Occupational Physiology 53 (53): 339–42. 10.1007/BF00422850.

Connick, Mark J., and François Xavier Li. 2013. “The Impact of Altered Task Mechanics on Timing and Duration of Eccentric Bi-Articular Muscle Contractions during Cycling.” Journal of Electromyography and Kinesiology 23 (23): 223–29. 10.1016/j.jelekin.2012.08.012.

Donelan, J. M., R. Kram, and A. D. Kuo. 2001. “Mechanical and Metabolic Determinants of the Preferred Step Width in Human Walking.” Proceedings of the Royal Society B: Biological Sciences 268 (268): 1985–92. 10.1098/rspb.2001.1761.

Elmer, Steven J., Paul R. Barratt, Thomas Korff, and James C. Martin. 2011. “Joint-Specific Power Production during Submaximal and Maximal Cycling.” Medicine and Science in Sports and Exercise 43 (43): 1940–47. 10.1249/MSS.0b013e31821b00c5.

Farina, Dario, Roberto Merletti, and Roger M. Enoka. 2014. “The Extraction of Neural Strategies from the Surface EMG: An Update.” Journal of Applied Physiology 117 (117): 1215–30. 10.1152/japplphysiol.00162.2014.

Fenn, Wallace. 1923. “A QUANTITATIVE COMPARISON BETWEEN THE ENERGY LIBERATED AND THE WORK PERFORMED BY THE ISOLATED SARTORIUS MUSCLE OF THE FROG.” The Journal of Physiology 58 (2–3): 111–258.

Fenn, Wallace. 1924. “The Relation between the Work Performed and the Energy Liberated in Muscular Contraction.” Journal of Physiology 58 (58): 373–95.

Gandevia, S. C. 2001. “Spinal and Supraspinal Factors in Human Muscle Fatigue.” Physiological Reviews 81 (81): 1725–89. 10.1152/physrev.2001.81.4.1725.

Griffin, Timothy M., Thomas J. Roberts, and Rodger Kram. 2003. “Metabolic Cost of Generating Muscular Force in Human Walking: Insights from Load-Carrying and Speed Experiments.” Journal of Applied Physiology 95 (95): 172–83. 10.1152/japplphysiol.00944.2002.

Handsfield, Geoffrey G., Craig H. Meyer, Joseph M. Hart, Mark F. Abel, and Silvia S. Blemker. 2014. “Relationships of 35 Lower Limb Muscles to Height and Body Mass Quantified Using MRI.” Journal of Biomechanics 47 (47): 631–38. 10.1016/j.jbiomech.2013.12.002.

Hermens, Hermie J, Bart Freriks, Catherine Disselhorst-Klug, and Günter Rau. 2000. “Development of Recommendations for SEMG Sensors and Sensor Placement Procedures.” Journal of Electromyography and Kinesiology 10 (10): 361–74.

Holt, Kenneth, Joseph Hamill, and Robert Andres. 1990. “Predicting the Minimal Energy Costs of Human Walking.”

Hoyt, Donald. 1981. “Gait and the Energetics of Locomotion in Horses.” Nature 292 (July): 239–40.

Kipp, Shalaya, William C. Byrnes, and Rodger Kram. 2018. “Calculating Metabolic Energy Expenditure across a Wide Range of Exercise Intensities: The Equation Matters” XXVIII: 1–14.

Kipp, Shalaya, Alena M. Grabowski, and Rodger Kram. 2018. “What Determines the Metabolic Cost of Human Running across a Wide Range of Velocities?” Journal of Experimental Biology 221 (221). 10.1242/jeb.184218.

Kram, Rodger, and C. Richard Taylor. 1990. “Energetics of Running: A New Perspective.” Nature 346 (346): 265–67. 10.1038/346265a0.

Kruschewsky, Alberto B., Rodolfo A. Dellagrana, Mateus Rossato, Luiz Fernando P. Ribeiro, Caetano D. Lazzari, and Fernando Diefenthaeler. 2018. “Saddle Height and Cadence Effects on the Physiological, Perceptual, and Affective Responses of Recreational Cyclists.” Perceptual and Motor Skills 125 (125): 923–38. 10.1177/0031512518786803.

Leavitt, Trevor G., and Heather K. Vincent. 2016. “Simple Seat Height Adjustment in Bike Fitting Can Reduce Injury Risk.” Current Sports Medicine Reports 15 (15): 130. 10.1249/JSR.0000000000000254.

MacIntosh, Brian R., Richard R. Neptune, and John F. Horton. 2000. “Cadence, Power, and Muscle Activation in Cycle Ergometry.” Medicine and Science in Sports and Exercise 32 (32): 1281–87. 10.1097/00005768-200007000-00015.

Marino, Frank E., Benjamin E. Sibson, and Daniel E. Lieberman. 2022. “The Evolution of Human Fatigue Resistance.” Journal of Comparative Physiology B: Biochemical, Systemic, and Environmental Physiology 192 (3–4): 411–22. 10.1007/s00360-022-01439-4.

Marsh, Anthony P., and Philip E. Martin. 1993. “The Association between Cycling Experience and Preferred and Most Economical Cadences.” MEDICINE & SCIENCE IN SPORTS & EXERCISE 2: 1269–74.

Marsh, Anthony P., and Philip E. Martin. 1997. “Effect of Cycling Experience, Aerobic Power, and Power Output on Preferred and Most Economical Cycling Cadences.” Medicine and Science in Sports and Exercise 29 (29): 1225–32. 10.1097/00005768-199709000-00016.

McDonald, Kirsty A., Joseph P. Cusumano, Andrew Hieronymi, and Jonas Rubenson. 2022. “Humans Trade off Whole-Body Energy Cost to Avoid Overburdening Muscles While Walking.” Proceedings of the Royal Society B: Biological Sciences 289 (1985). 10.1098/rspb.2022.1189.

Miller, Ross H., Brian R. Umberger, Joseph Hamill, and Graham E. Caldwell. 2012. “Evaluation of the Minimum Energy Hypothesis and Other Potential Optimality Criteria for Human Running.” Proceedings of the Royal Society B: Biological Sciences 279 (279): 1498–1505. 10.1098/rspb.2011.2015.

Minetti, A. E., and R. Mc N. Alexander. 1997. “A Theory of Metabolic Costs for Bipedal Gaits.” Journal of Theoretical Biology 186 (186): 467–76. 10.1006/jtbi.1997.0407.

Monjo, Florian, Romain Terrier, and Nicolas Forestier. 2015. “Muscle Fatigue as an Investigative Tool in Motor Control: A Review with New Insights on Internal Models and Posture-Movement Coordination.” Human Movement Science 44: 225–33. 10.1016/j.humov.2015.09.006.

Neptune, R. R., and M. L. Hull. 1999. “A Theoretical Analysis of Preferred Pedaling Rate Selection in Endurance Cycling.” Journal of Biomechanics 32 (32): 409–15. 10.1016/S0021-9290(98)00182-1.

Nordeen-Snyder, Katherine. 1977. “The Effect of Bicycle Seat Height Variation upon Oxygen Consumption and Lower Limb Kinematics.”

Peveler, Will. 2008. “Effects of Saddle Height on Economy in Cycling.” Journal of Strength and Conditioning Research 22 (22): 1355–59.

Pontzer, Herman. 2017. “Economy and Endurance in Human Evolution.” Current Biology 27 (27): R613–21. 10.1016/j.cub.2017.05.031.

Proske, Uwe, and Trevor Allen. 2019. “The Neural Basis of the Senses of Effort, Force and Heaviness.” Experimental Brain Research 237 (237): 589–99. 10.1007/s00221-018-5460-7.

Ralston, H. J. 1958. “Energy-Speed Relation and Optimal Speed during Level Walking.” Internationale Zeitschrift Für Angewandte Physiologie Einschliesslich Arbeitsphysiologie 17 (17): 277–83. 10.1007/BF00698754.

Rankin, Jeffery W., and Richard R. Neptune. 2010. “The Influence of Seat Configuration on Maximal Average Crank Power during Pedaling: A Simulation Study.” Journal of Applied Biomechanics 26 (26): 493–500. 10.1123/jab.26.4.493.

Rasmussen, John, Michael Damsgaard, and Michael Voigt. 2001. “Muscle Recruitment by the Min/Max Criterion - A Comparative Numerical Study.” Journal of Biomechanics 34 (34): 409–15. 10.1016/S0021-9290(00)00191-3.

Riveros-Matthey, Cristian, Timothy J. Carroll, Mark J. Connick, and Glen A. Lichtwark. 2023. “An In-Silico Investigation of the Effect of Changing Cycling Crank Power and Cadence on Muscle Energetics and Active Muscle Volume.” BioRxiv C: 1–29.

Riveros-Matthey, Cristian, Timothy J. Carroll, Glen A. Lichtwark, and Mark J. Connick. 2023. “The Effects of Crank Power and Cadence on Muscle Fascicle Shortening Velocities, Muscle Activations and Joint-Powers during Cycling.” Journal of Experimental Biology. 10.1242/jeb.245600.

Riveros-Matthey, Cristian D., Mark J. Connick, Glen A. Lichtwark, and Timothy J. Carroll. 2024. “Saddle Height Variation: Cycling Muscle Activation & Energy Data.” UQ ESpace. 10.48610/6413ffe.

Rugg, S.G., and R.J. Gregor. 1987. “THE EFFECT OF SEAT HEIGHT ON MUSCLE LENGTHS, VELOCITIES AND MOMENT AIL”LIE NGTHS DURING CYCLING,” 899.

Russell, Daniel M., and Dylan T. Apatoczky. 2016. “Walking at the Preferred Stride Frequency Minimizes Muscle Activity.” Gait and Posture 45: 181–86. 10.1016/j.gaitpost.2016.01.027.

Sanderson, David J., and Annita T. Amoroso. 2009. “The Influence of Seat Height on the Mechanical Function of the Triceps Surae Muscles during Steady-Rate Cycling.” Journal of Electromyography and Kinesiology 19 (19): 465–71. 10.1016/j.jelekin.2008.09.011.

Selinger, Jessica C., Shawn M. O’Connor, Jeremy D. Wong, and J. Maxwell Donelan. 2015. “Humans Can Continuously Optimize Energetic Cost during Walking.” Current Biology 25 (25): 2452–56. 10.1016/j.cub.2015.08.016.

Selinger, Jessica C., Jeremy D. Wong, Surabhi N. Simha, and J. Maxwell Donelan. 2019. “How Humans Initiate Energy Optimization and Converge on Their Optimal Gaits.” Journal of Experimental Biology 222 (222). 10.1242/jeb.198234.

Silder, Amy, Thor Besier, and Scott L. Delp. 2012. “Predicting the Metabolic Cost of Incline Walking from Muscle Activity and Walking Mechanics.” Journal of Biomechanics 45 (45): 1842–49. 10.1016/j.jbiomech.2012.03.032.

Snaterse, Mark, Robert Ton, Arthur D. Kuo, and J. Maxwell Donelan. 2011. “Distinct Fast and Slow Processes Contribute to the Selection of Preferred Step Frequency during Human Walking.” Journal of Applied Physiology 110 (110): 1682–90. 10.1152/japplphysiol.00536.2010.

Sousa, Andreia S.P., and João Manuel R.S. Tavares. 2012. “Effect of Gait Speed on Muscle Activity Patterns and Magnitude during Stance.” Motor Control 16 (16): 480–92. 10.1123/mcj.16.4.480.

Takaishi, Tetsuo, Yoshifumi Yasuda, Takashi Ono, and Toshio Moritani. 1996. “Optimal Pedaling Rate Estimated from Neuromuscular Fatigue for Cyclists.” Medicine and Science in Sports and Exercise 28 (28): 1492–97. 10.1097/00005768-199612000-00008.

Umberger, Brian R. 2010. “Stance and Swing Phase Costs in Human Walking.” Journal of the Royal Society Interface 7 (7): 1329–40. 10.1098/rsif.2010.0084.

Umberger, Brian R., and Philip E. Martin. 2007. “Mechanical Power and Efficiency of Level Walking with Different Stride Rates.” Journal of Experimental Biology 210 (210): 3255–65. 10.1242/jeb.000950.

Wolpert, D. M., and J. R. Flanagan. 2001. “Motor Prediction.” Current BiologylJ: CB 11 (11): 729–32. 10.1016/s0960-9822(01)00432-8.

Wong, Jeremy D., Jessica C. Selinger, and J. Maxwell Donelan. 2019. “Is Natural Variability in Gait Sufficient to Initiate Spontaneous Energy Optimization in Human Walking?” Journal of Neurophysiology 121 (121): 1848–55. 10.1152/jn.00417.2018.

Zarrugh, M. Y., F. N. Todd, and H. J. Ralston. 1974. “Optimization of Energy Expenditure during Level Walking.” European Journal of Applied Physiology and Occupational Physiology 33 (33): 293–306. 10.1007/BF00430237.

